# Environmental variability and the evolution of bet-hedging strategies

**DOI:** 10.1101/2020.02.29.971432

**Authors:** Kelley Slimon, Rachel M. Germain

**Affiliations:** Department of Zoology and the Biodiversity Research Centre, University of British Columbia, Vancouver, British Columbia, Canada

**Keywords:** adaptation, bet hedging, dispersal, dormancy, environmental variability, McLaughlin Natural Reserve

## Abstract

Bet-hedging strategies, such as dispersal and dormancy, are predicted to evolve in varying and uncertain environments and are critical to ecological models of biodiversity maintenance. Theories of the specific ecological scenarios that favor the evolution of dispersal, dormancy, or their covariance are rarely tested, particularly for naturally-evolved populations that experience complex patterns of spatiotemporal environmental variation. We grew 23 populations of *Vulpia microstachys*, an annual grass native to California, in a greenhouse, and on the offspring generation measured seed dispersal ability and dormancy rates. We hypothesized that seed dormancy rates and dispersal abilities would be highest in populations from more productive, temporally variable sites, causing them to covary positively. Contrary to our hypothesis, our data suggest that both dispersal and dormancy evolve to combat different axes and scales of spatial heterogeneity, and are underlain by different seed traits, allowing them to evolve independently. Dormancy appears to have evolved as a strategy for overcoming microgeographic heterogeneity rather than temporal climate fluctuations, an outcome that to our knowledge has not been considered by theory. In sum, we provide much needed empirical data on the evolution of bet hedging, as well as a new perspective on the ecological function dormancy provides in heterogeneous landscapes.

## Introduction

Most species in nature contend with variable, uncertain conditions in some portion of their geographic range (Vasseur and Yodzis 2004; Coumou and Rahmstorf 2012) posing a challenge to adaptive evolution (Bell 2010). In variable environments, an adaptive response to conditions at a given point in time might be maladaptive on longer timescales, for example, if the direction of selection fluctuates inter-annually (Hamann et al. 2018). Although phenotypic plasticity, i.e., the ability of a single genotype to produce different phenotypes in different environments, might evolve in response to frequent and predictable variation in environmental conditions, it may be selected against if fluctuating conditions are unpredictable (Simons 2011), such as when fluctuations are infrequent and extreme (Chevin and Hoffmann 2017) or sample a great range of multidimensional environmental space. These difficult-to-adapt-to scenarios are exactly the scenarios that are expected to increase in frequency with climate change (Easterling et al. 2000), such as the incidence of extreme, unpredictable events (e.g., hurricanes (Dale et al. 2001), climate anomalies (Mahony et al. 2017)).

To combat environmental uncertainty when fluctuations are unpredictable, populations evolve bet-hedging strategies (Simons 2011), allowing some proportion of individuals to avoid or escape unsuitable conditions (Haaland et al. 2019). Across the tree of life, dispersal and dormancy are two common forms of bet hedging. The first strategy, dispersal, is the movement of individuals among populations, and is predicted to evolve in environments with low spatial and temporal autocorrelation (Venable and Brown 1988). The second strategy, dormancy, is a reversible state of reduced metabolic activity that can last for multiple generations, allowing individuals to persist through unfavourable periods. Dormancy is predicted to evolve when temporal autocorrelation is low but spatial autocorrelation is high (Venable and Brown 1988). In the absence of variability (i.e., high autocorrelation in space and time), neither strategy is predicted to evolve given that, by definition, bet-hedging strategies are costly to maintain as they require some individuals to forgo fitness even in sites and years that are suitable (i.e., by dispersing elsewhere or remaining dormant (Venable 2007; Siewert and Tielbörger 2010)). Unlike phenotypic plasticity, which in unpredictable environments, would require an individual to possess all the genetic machinery to express many phenotypes to combat many possible environmental challenges, bet hedging strategies instead allow individuals to tolerate or escape unsuitable conditions entirely (Simons 2011). Note that bet hedging strategies, such as dormancy and dispersal, can be plastic (Rees et al. 2010; Maxwell and Magwene 2017), but plasticity itself is not a form of bet hedging (Joschinski and Bonte 2019).

Although theory of when dispersal and dormancy evolve appears straightforward, two factors complicate predictions in real systems. First, in environments that experience variation both in space and time, theory differs in whether dispersal, dormancy, or some combination of both will evolve (Casas et al. 2015). As both dispersal and dormancy are forms of bet hedging, each of which has associated fitness costs, early models by Cohen and Levin (1987) and Venable and Brown (Venable and Brown 1988) emphasize that their evolution is constrained by a tradeoff, causing negative covariance. By contrast, more recent models predict positive covariance due to genetic linkages or pleiotropy (Wisnoski et al. 2019) or when environmental conditions are strongly autocorrelated in both space and time (Snyder 2006). Dispersal and dormancy can also evolve independently if each is underlain by distinct, genetically-unlinked traits (Arnold 1992). Despite a large body of theory, few empirical studies to our knowledge quantify dispersal-dormancy covariance (e.g., (Rees 1993; Siewert and Tielbörger 2010)). General support for any specific theoretical prediction is not yet possible, limiting our understanding of adaptation in variable environments (Casas et al. 2015).

Second, predictions of when dispersal and dormancy evolve are based on the magnitude of autocorrelation in space and time, which contrasts the complex structure of environmental variance in nature. Due to interactions between spatial landscape heterogeneity and temporal climate variability, temporal variability is often not realized equally across space. In grassland ecosystems, for example, plant biomass varies by orders of magnitude among years in response to climate fluctuations, but this effect is greatly dampened in unproductive environments (Fig. 1, yellow line (Eskelinen and Harrison 2015)). Individuals existing in locations of a landscape that are unproductive relative to other locations tend to be more limited by local resource conditions than by climate, meaning that even in years that are climatically favorable, individuals lack the materials to respond—this phenomenon is known as ‘co-limitation’ (Eskelinen and Harrison 2015). In productive locations on a landscape, local resources are not as limiting, leading to communities which produce higher biomass on average but which also experience high biomass variability as climates fluctuate (Fig. 1, orange line). As a consequence of co-limitation, the evolution of dispersal and dormancy may vary spatially, even among populations within a single region—evolving more commonly in habitat patches that are more suitable on average but that also experience high variability. Note that co-limitation is likely a general phenomenon beyond plants, and presents a unique ecological problem for evolution to solve that has not been considered by existing theory.

**Figure 1.**
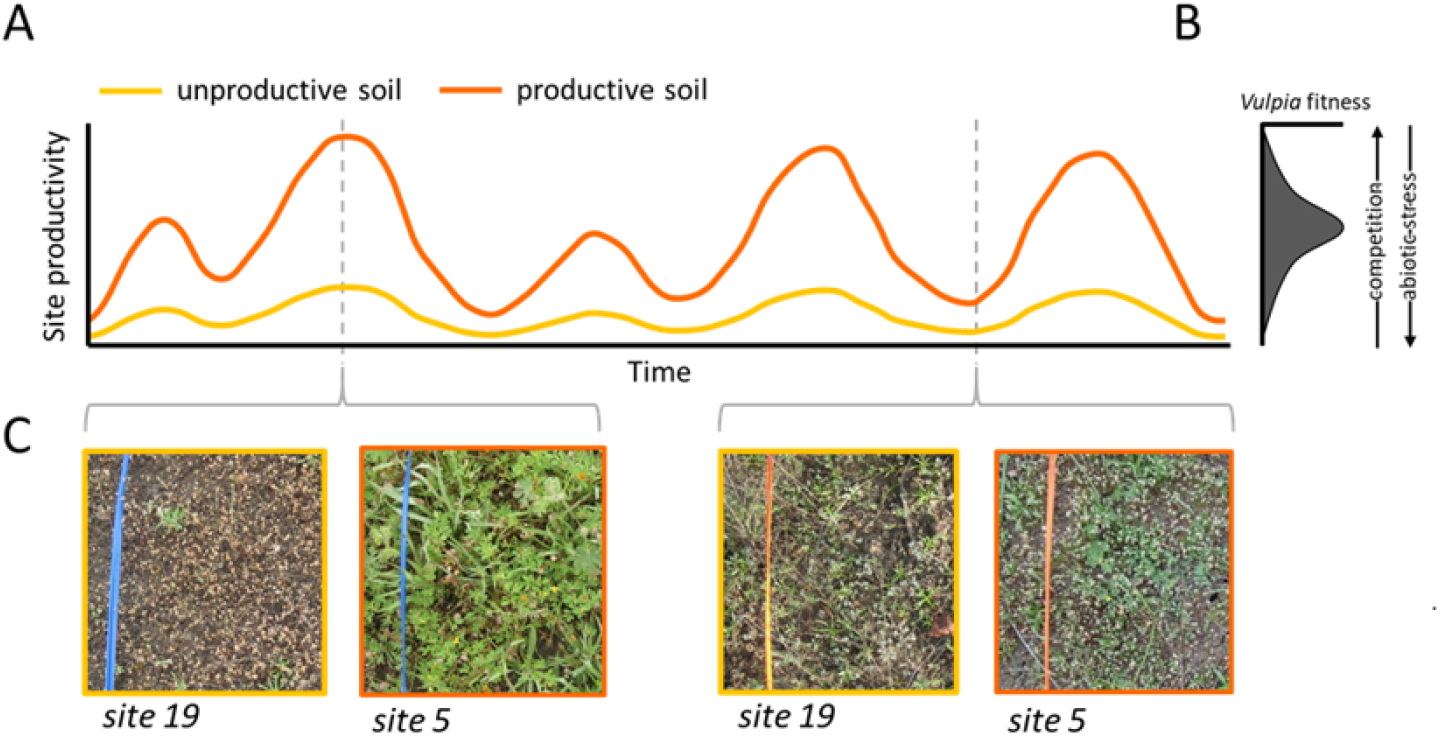
Schematic illustrating the phenomenon of ‘co-limitation’, which causes variability in plant productivity due to climate to be dampened in sites from unproductive soils (A). Dormancy rates but not dispersal ability varied along a site productivity gradient (A (i) and B (i)) whereas dispersal ability but not dormancy rates varied with site pH and soil moisture content (A (ii) and B (ii)). Points are the fitted population averages from a glme model (orange lines showing regression fits), with grey boxplots showing the distribution of the raw data within populations. The surface in the 3d figures are the fitted relationships from glme models.

We tested how dispersal, dormancy, and their covariance have evolved in 23 natural populations of an annual grass (*Vulpia microstachys*) along environmental gradients. *Vulpia microstachys* is native to North America and has a seed morphology known to confer dispersal by attachment to passing animals (Pijl and van der Pijl 1972; Rees 1993). At our study site in Northern California, *V. microstachys* is found in 87% of surveyed serpentine habitat, spanning a broad gradient of environmental conditions (Germain et al. 2017). Co-limitation has been shown to occur in serpentine grassland communities at our study site in past research (Eskelinen and Harrison 2015), demonstrating that plants in harsh serpentine habitat patches experience less variability than in benign patches (also see Fig. 1C). Additionally, previous research with serpentine populations of *V. microstachys* demonstrates that it has highest fitness at intermediate productivities, specifically when abiotic conditions are not too harsh but competition is not yet intense (Fig. 1b (Jurjavcic, Harrison, and Wolf, 2002)). As bet-hedging strategies serve to cope with environmental variability, we predicted that dispersal, dormancy, or both would have evolved to be higher in populations from more productive patches, since interannual climate fluctuations cause conditions to vary in suitability through time (orange line in Fig. 1B). To test these predictions, we propagated seeds of field-collected *V. microstachys* populations in a common environment to standardize the maternal environment, and on the offspring generation, measured seed dormancy and seed dispersal ability. As we will discuss, our predictions were not supported. Dispersal and dormancy both evolved but did not covary, and the evolution of dormancy countered theory but can be understood by examining the spatial scale of environmental heterogeneity.

## Methods

### Study system

Our experiments included 23 populations of *Vulpia microstachys* collected from serpentine habitat at the UC McLaughlin Natural Reserve in Lower Lake, CA (38.873889N, 122.431667W). Serpentine soils are common in subduction zones, such as along the San Andreas fault, where the Earth’s mantle becomes exposed. Serpentine soils are characterized by high Mg/Ca ratios, high heavy metal content, low soil fertility, and often a rocky texture (Jurjavcic et al. 2002). Together, these variables correlate with overall plant productivity and diversity, with “harsh” sites characterized by low productivity, low diversity, and a nested subset of species that can tolerate harsh conditions (including *V. microstachys* (Armstrong et al. 1992)). Additionally, serpentine habitat is highly fragmented, both by an unsuitable nonserpentine matrix dominated by Avena spp. and by other intervening habitat, such as oak woodlands.

### Seed collection and maternal generation

For each population, seeds were bulk-collected from hundreds of plants in October 2016, before fall rains and following summer heat-stratification, and grown at the research greenhouses at the University of British Columbia. Before any trait data were collected, we grew the field-collected seeds for one generation to remove the potential for maternal environmental effects to obfuscate signals of evolution. For the maternal generation, seeds of each population were propagated in 10 replicate 2.79 L pots, with each pot containing approximately 50 seeds. There were 230 pots in total (23 populations x 10 replicate pots) arranged on greenhouse benches in a completely randomized design. Growing conditions were set to mimic the progression of a growing season typical of Lower Lake, CA. The greenhouse was initially set to mimic cool, wet winter conditions (15/7C day/night cycle with 11 hours of daylight, supplemented by high intensity discharge lighting). As the growing season progressed, gradually over six months, we increased temperatures to 30/15C and a day length of 16 hours. The pots were top-watered to keep the soil constantly moist as plants established, and were then bottom-watered as needed after establishment (at an interval that allowed the soil to dry out completely). Fertilizer was added twice and powdery mildew was spot treated with MilStop fungicide as needed. Because the goal of the maternal generation was propagation of seeds to be used in future experiments, we used a standard high fertility potting soil (Sunshine Mix 5). Seeds were collected as they matured, dried at 60C for 72 hours, and stored in coin envelopes.

### Offspring generation

Seeds produced in the maternal generation were used in two separate experiments, both initiated September 2017 on adjacent greenhouse benches: one to measure seed dormancy rates and the other seed dispersal ability. For the dormancy experiment, we filled 2.54 L pots with a 1:4 ratio of Sunshine potting mix and serpentine soil collected from Grasshopper Mountain in British Columbia (4931’57.10”N, 12054’12.94”W). Grasshopper Mountain is biochemically similar to McLaughlin serpentine. We did not use soil from McLaughlin because we wanted a soil to which all populations were naïve, as soil biota can complicate evolutionary signals (Aarssen and Turkington 1985). Seven seeds of *V. microstachys* were added to each pot, with 10 replicate pots per each of the 23 populations. Pots were arranged in a completely randomized design on a greenhouse bench under climate conditions similar to the maternal generation. We tracked what proportion of seeds per pot germinated over a six week period, estimating dormancy rates as the number of ungerminated seed.

For the dispersal ability experiment, seeds were sewn into clear 2.25” 10” pots, suspended at a 45 angle and covered with a thin layer of peat moss. This specialized pot was used to perform non-destructive root measurements as part of an unrelated trait experiment. For each population there were 10 replicate pots each containing seven seeds, arranged on greenhouse benches in a completely randomized design. We tracked germination success and thinned densities to a single randomly-selected individual. All seeds were collected from each plant as they senesced (separately per pot), dried at 60C for 72 hours, and stored in coin envelopes. To estimate dispersal ability, 20 seeds from each envelope (i.e., per pot) were laid out in a grid on a lab bench. A deer “leg” (a deer pelt fixed to a cylinder) was systematically rolled over each grid of seeds, which was then dropped down a dowel from a 15-cm height onto a rubber base to simulate the downward force of walking. The number of seeds to attach initially and that remained attached after the force was applied was recorded. This method of testing dispersal of vertebrate-dispersed plant species using simulated animal limbs has been employed by past studies (Carlquist and Pauly 1985; Mouissie et al. 2005).

#### Stats

To test whether population-level dormancy and dispersal rates varied as a function of the productivity of the sites they originated from, we ran generalized linear mixed effects (glme) models with a binomial error distribution using R package ‘glmmTMB’ (v0.2.3 (Brooks et al. 2017)). The number of seeds that germinated or attached to a deer pelt was the response variable, the total number of seeds included in the trial was included as a weight, site productivity was a fixed effect, and population was a random effect. The dormancy results presented in the main text were performed in a harsh germination environment, as dormancy rates were very high compared to those of a separate concurrent germination trial (Fig. S1)—these trials ran concurrently but differed in whether or not seeds were protected by a thin layer of peat moss. This difference tells us that dormancy responds plastically to germination cues, and that populations from harsher environments have evolved seeds with an increased ability to sense and respond to those cues.

As we will discuss, site productivity did not significantly predict dispersal rates, but including ‘population’ as a random effect significantly improved model fit in comparison to an intercept-only model. To explain observed differences in dispersal ability among populations, we borrowed existing data of eight environmental variables (% soil moisture, N, P, K, Na, cation exchange capacity, pH, and Mg/Ca ratios) available for 20 of our populations from previous research (Germain et al. 2017). We repeated the glme models above, except included the eight environmental variables as additive fixed effects instead of productivity. We performed backwards model selection using the ‘step’ function until a reduced model with the lowest AIC scores was obtained. This final reduced model included % soil moisture and pH; we present these results as a 3d surface generated using the ‘visreg2d’ function in package ‘visreg’(v2.6-0 (Breheny and Burchett 2017)).

To examine if dormancy rates and dispersal ability significantly covaried, we used R package ‘lmodel2’ to perform a major axis regression permutation test (v1.7-3 (Legendre 1998)). Major axis regression is appropriate when two variables are dependent on one another, are each measured with error, and covary linearly. This contrasts regular simple linear regression where the assumption is that one variable is independent and one is dependent. Each variable was a vector of fitted population averages from our glme models. We used 999 permutations.

### Seed morphology

The dormancy and dispersal measurements described above represent organismal functions that are underlain by specific traits, such as morphology and physiology (Losos 2011). In order for dormancy and dispersal to evolve independently, they must be underlain by different traits that each have a genetic basis (Arnold 1992). Very little is known about which specific traits affect dormancy and dispersal in *V. microstachys*, but heavier seeds with long awns are suggested to have higher germination rates (Peart 1981; Garnier and Dajoz 2001; Venable 2007) whereas seeds with long awns and long hairs are hypothesized to attach more readily to passing animals (Pijl and van der Pijl 1972; Rees 1993).

We measured seed mass, seed awn length, and seed hair length on the same seed samples used in the dispersal ability trials. We measured seed mass by weighing five replicate samples of 50 seed per population, dividing each sample by 50 to estimate mean mass per seed. Awn length was measured by electronic caliper to the nearest 0.01 mm, and seed hair length was a measurement of the longest hair present on each seed to the nearest 0.005 mm taken using a dissecting scope mounted with a stage micrometer. Awn length and seed hair length were both measured on five seeds chosen at random from each sample envelope, which we averaged.

#### Stats

We used separate glme models with identical model structure to test if seed traits determined the numbers of seeds that were dormant and that attached to the deer pelt. We used a binomial error distribution with the total number of seeds included in the trial as a weight, so that proportion successes is adjusted by the total number of trials. We used seed mass, seed awn length, and seed hair length as additive fixed effects and population as a random effect.

### Spatial scale of environmental suitability

We found that dormancy rates decreased with site productivity, a counterintuitive result given that unproductive sites experience less temporal variability. An alternative explanation for this result is that fine-scale spatial heterogeneity in microsite suitability is greater in unproductive sites, for example, if they were rockier and less penetrable to plant roots. Dormancy would thus act as a mechanism allowing short-range dispersal if a seed happened to be dropped in an unsuitable microsite (Wisnoski et al. 2019). To test this alternative explanation, we returned to twelve sites at McLaughlin Natural Reserve in May 2019, six of the highest productivity sites and six of the lowest productivity. At each site, we haphazardly selected five plots by throwing a 25 × 25 cm quadrat 10 m in random directions, and within each plot, created a grid of 25 microsites. At each microsite, we tested if a six-gauge nail could penetrate the ground by at least 0.5 inches, recording successes as a potential suitable microsite and failures as unsuitable microsites. We consider this estimation of microsite suitability as highly conservative as some penetrable microsites were devoid of vegetation in the unproductive sites.

#### Stats

We analyzed these data with a mixed effects logistic regression with site productivity as the explanatory variable (low vs. high), microsite suitability as the binary response variable, and site as a random effect to account for multiple plots per site. The model was weighted by the total number of trials in a plot (i.e., 25).

## Results

We predicted that dormancy rates would have evolved to be greatest in productive environments, given that productive environments vary more strongly with climate than unproductive sites (Eskelinen and Harrison 2015)—we instead found the opposite pattern (Fig. 2A; χ2 = 19.9, P *<* 0.001). Populations from the least productive sites produced seed with predicted dormancy rates of 80% compared to 54% in the most productive sites. An explanation for this counterintuitive result might be that unproductive sites are more variable in fine-scale heterogeneity than productive sites, owing to more rocky, coarse substrates. This would select for the evolution of increased dormancy if seeds were frequently deposited in unsuitable microsites and remained dormant until rain, wind, or disturbance moved them short distances (*<*25cm) to a suitable microsite. To test this new, alternative hypothesis, we returned to the field in 2019 to test for differences in microsite suitability along the productivity gradient (see Methods), and indeed found that unproductive sites contained 3.25x more unsuitable microsites than productive sites at a spatial scale relevant to local seed movement (Fig. 3; χ2 = 16.6, P *<* 0.001).

**Figure 2.**
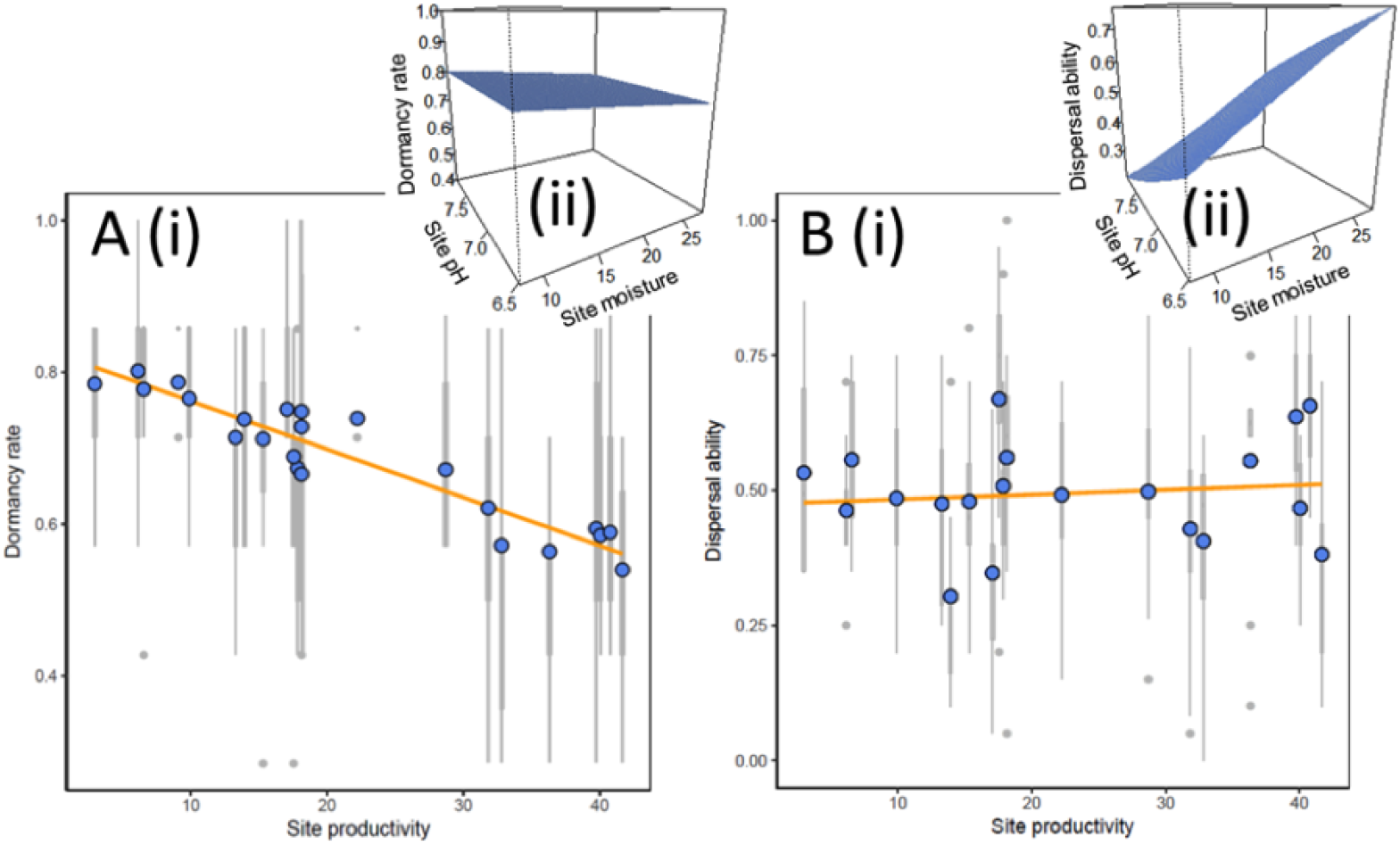
Environmental drivers of population differentiation in dormancy rates (A) and dispersal ability (B). Dormancy rates but not dispersal ability varied along a site productivity gradient (A (i) and B (i)) whereas dispersal ability but not dormancy rates varied with site pH and soil moisture content (A (ii) and B (ii)). Points are the fitted population averages from a glme model (orange lines showing regression fits), with grey boxplots showing the distribution of the raw data within populations. The surface in the 3d figures are the fitted relationships from glme models.

**Figure 3.**
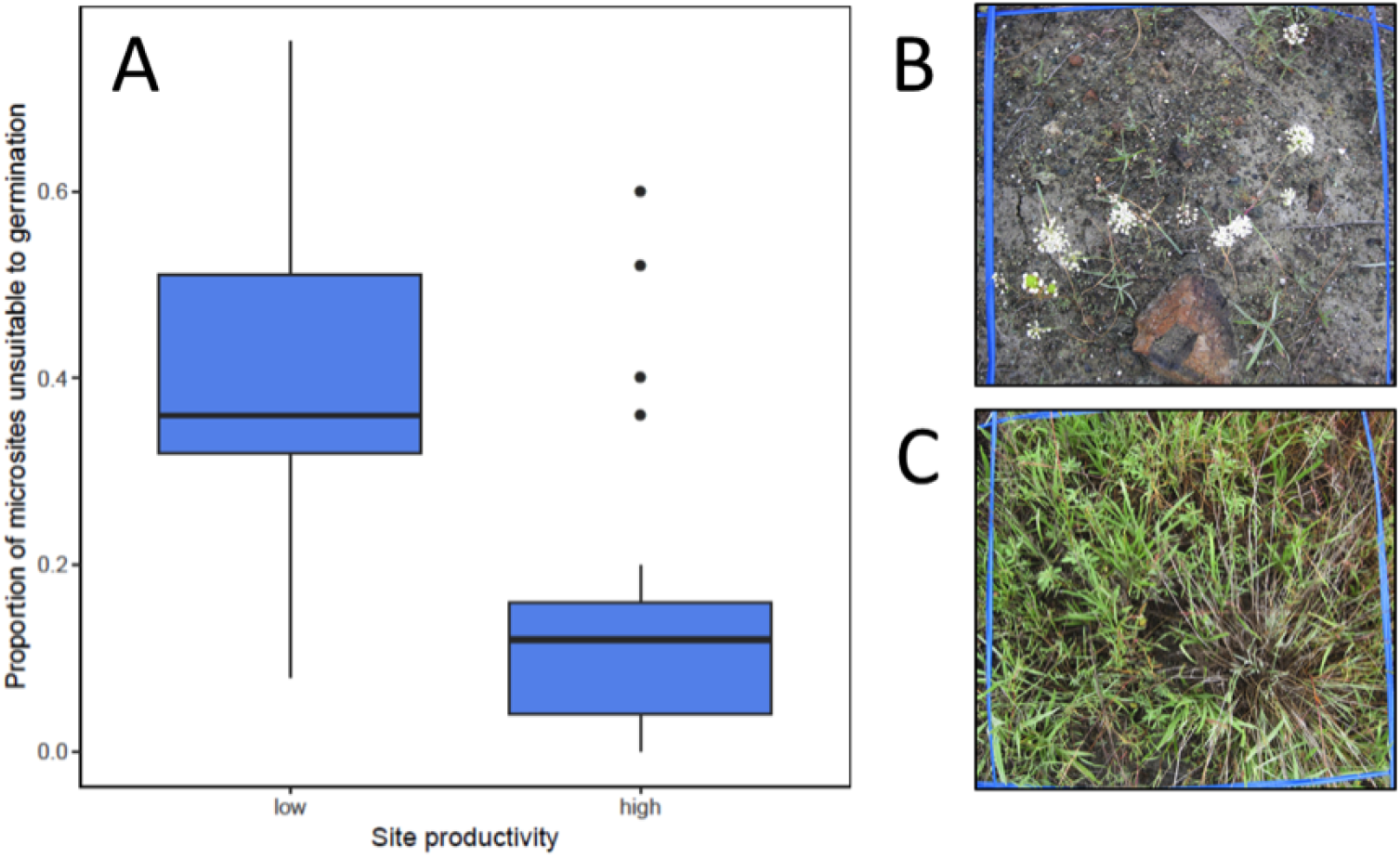
Microsite limitation is greater in low productivity sites (A), shown as boxplots. Low productivity sites contain more rocks and other substrates impenetrable to plant roots, leading to sparse vegetation (B) compared to high productivity sites (C)—we show one site per treatment group as examples. n = six sites per treatment, five plots per site, 25 microsites tested per plot; outlying points are individual plots.

In contrast to dormancy rates, we predicted that dispersal ability would have evolved to be highest in high productivity sites and instead found no relationship (Fig. 2B; χ2 = 0.2, P = 0.659). On average, populations exhibited 50% adherence to deer fur regardless of productivity, ranging from 30% to 67% adherence. Although this result could indicate that dispersal has not evolved, a comparison among models with and without population as a random effect demonstrated that population explained a significant amount of variation in dispersal ability (AICc = 110.6; P *<* 0.001). We used model selection of additional environmental variables to pinpoint alternative environmental drivers of population differentiation, which identified a model which included significant additive effects of soil moisture (χ2 = 16.2, P *<* 0.001) and pH (χ2 = 11.3, P *<* 0.001). Specifically, populations from sites with high soil moisture and low pH had the highest dispersal ability (Fig. 2B (ii)). We tested if dormancy rates also varied as a function of site pH and soil moisture and did not find statistically significant relationships (Fig. 2A (ii); both P *>* 0.50).

Despite the fact that dispersal and dormancy both function as bet-hedging strategies, suggesting that they may evolve in tandem (positive covariance) or be constrained by a tradeoff (negative covariance), we found that dispersal and dormancy did not covary with each other (Fig. 4; major axis regression: r =−0.096, two-tailed P = 0.687). We can understand this result by considering two pieces of evidence. First, we have already shown that dispersal and dormancy have evolved along different environmental gradients (Fig. 2) meaning that the assumption that both strategies are equal and alternative forms of bet hedging may not be correct (Husband and Barrett 1996). Second, we examined three morphological seed traits posed in the literature as conferring dispersal, dormancy, or both: seed mass, awn length, and hair length. We found that dispersal and dormancy were not underlain by the same traits (Table 1). Specifically, dormancy rate was not correlated with any seed trait despite previous research suggesting that seeds that are large in size or that have long awns are less dormant (Peart 1981; Garnier and Dajoz 2001; Venable 2007) whereas seeds with longer hairs had increased dispersal ability, attaching and holding on to deer fur at a higher rate than seeds with short hairs (Table 1), consistent with past research (Pijl and van der Pijl 1972; Rees 1993). A lack of covariance is possible when dispersal and dormancy are underlain by distinct traits and evolve in response to different environmental variables (Wisnoski et al. 2019), as appears to be the case in our study.

**Table 1.** The effect of morphological traits of seeds on dormancy rates, initial rate of attachment to a deer leg, rate of continued attachment after walking force is applied, and the total proportion adherence.

|  | Awn length |  | Hair length |  | Seed size |  |
| --- | --- | --- | --- | --- | --- | --- |
|  | Estimate | *P* | Estimate | *P* | Estimate | *P* |
| ***Dormancy*** | -0.15 | 0.162 | -0.01 | 0.706 | 0.12 | 0.189 |
| ***Dispersal*** |  |  |  |  |  |  |
| Initial attachment | -0.04 | 0.713 | -0.21 | **0.003** | 0.13 | 0.695 |
| Attachment after force applied | -0.05 | 0.157 | -0.09 | **<0.001** | 0.20 | 0.077 |
| Total attachment | -0.04 | 0.292 | -0.15 | **<0.001** | 0.13 | 0.251 |

**Figure 4.**
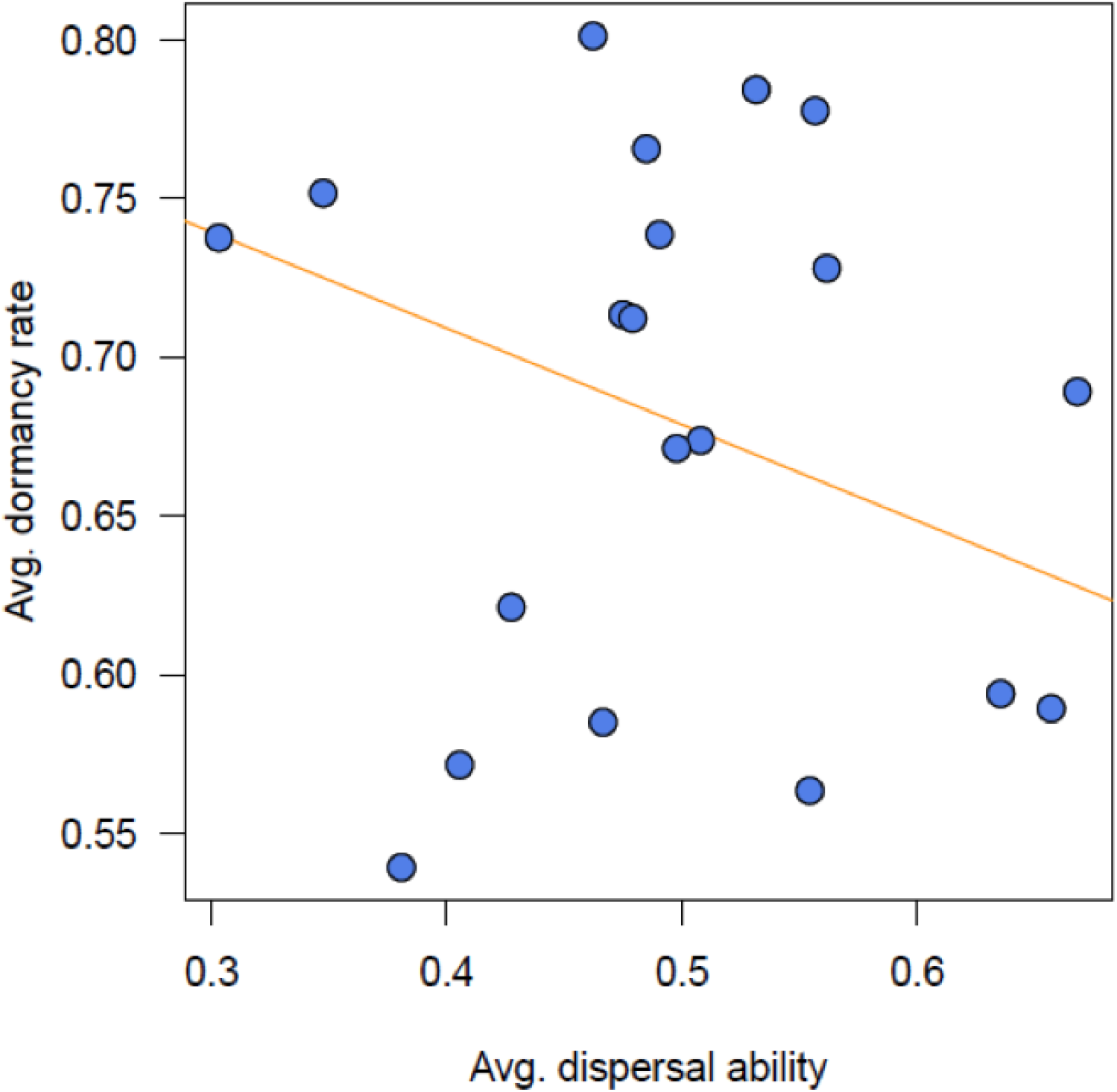
Dormancy rates and dispersal abilities do not covary. Points are population-level averages fitted from glme models and the orange line is the fitted relationship from a major axis regression (r = −0.096, two-tailed P = 0.687).

## Discussion

The structure of environmental variability in space, time, and their intersection is complex in most landscapes (Bell and Lechowicz 1991; Vasseur and Yodzis 2004; Fernandez-Going et al. 2013; Eskelinen and Harrison 2015), complicating predictions of how bet-hedging strategies evolve (Casas et al. 2015). Consistent with this idea, we show that different sources and scales of variability have selected for different types of bet-hedging strategies, allowing dispersal and dormancy to evolve independently. Below, we weigh our findings against relevant evolutionary theories. As adaptive evolution can be viewed as nature finding a solution to an ecological problem, we will also discuss how our results elucidate the ecological value of dormancy, dispersal, and their interaction in populations and metapopulations.

Few studies have examined the relationship between dormancy and dispersal in natural populations (Rees 1993; Siewert and Tielbörger 2010), highlighting the need for more studies, such as ours. Of those few studies, one reported a negative relationship (Rees 1993) while others found that dispersal and dormancy varied independently (Siewert and Tielbörger 2010). In Siewert and Tielbörger (2010), the authors posit that dispersal does not likely function as a bet hedging strategy in their desert annual system (i.e., there are no fitness benefits to dispersing), resulting in the lack of a relationship between dormancy and dispersal. Although we also found no relationship between dormancy and dispersal (Figure 4), the lack of dormancy-dispersal covariance we observed can be explained by each experiencing distinct agents and targets of selection rather than a lack of benefits (Wadgymar et al. 2017)—dispersal ability evolves in sites with low pH and high soil moisture content (the agent of selection) and is underlain by seed hair length (the likely target of selection) whereas dormancy rates evolve in unproductive sites and are underlain by some unmeasured (possibly physiological (Baskin and Baskin 2004)) trait. Similarly to our study and unlike Siewert and Tielbörger (2010), Venable (1998) found that dispersal and dormancy of Heterosperma pinnatum populations evolved in response to distinct environmental axes (vegetation for the former, precipitation for the latter). What is clear is that dormancy and dispersal are not simply alternative solutions, but solve distinct ecological problems

Counterintuitively, we found that seed dormancy was highest in populations from unproductive, less temporally variable environments. This observation would have run counter to theory had we not collected additional data showing that unproductive environments also had fewer suitable microsites. A recent perspective paper argues that dormancy may actually facilitate dispersal through unsuitable habitat (Wisnoski et al. 2019), linking habitat patches on regional scales. Our data suggests dormancy might be critical to combating small scale environmental heterogeneity, allowing seeds the opportunity to move short distances if first deposited in an unsuitable microsite. This fine spatial scale at which dormancy is under selection might help explain why at large scales, such as biogeographic scales, past work shows that dormancy does not evolve along climate gradients (Mayfield et al. 2014). Note that in our study populations from high productivity sites still had some baseline level of seed dormancy, with 50% of seeds remaining dormant in suboptimal germination conditions as opposed to 5% under optimal conditions (Fig. S1). In other words, populations from productive sites still enter dormancy but to a lesser extent than populations from unproductive sites.

Our finding that increased dispersal ability has evolved in certain environments suggests that gene flow among populations might occur asymmetrically in a landscape. Important to this argument is the certitude that dispersal does not result in gene flow if dispersing individuals fail to establish. If individuals do establish, and do so frequently, we would expect the recipient population to evolve increased dispersal ability as individuals that successfully disperse are more likely to express traits that confer increased dispersal ability—this is a key feature of spatial sorting during range expansions (Ochocki and Miller 2017) which also applies to metapopulations (Travis and Dytham 1998). However, in order for this outcome to occur, there must be an asymmetry in which types of environments dispersing individuals are likely to establish in. With knowledge of the natural history of our specific study system, we propose two explanations why establishment may be unequal across our landscape: (1) differences in habitat quality (i.e., in terms of average fitness value (Fig. 1B)), which affect both establishment success and the absolute number of dispersers that leave a patch. In studies of local adaptation, it is commonly observed that populations from both poor and high quality sites perform better in high quality sites, even if individuals from poor sites have adaptations to cope with harsh conditions ((Kawecki and Ebert 2004); e.g., dormancy, as shown here). As a consequence, in our study system, establishment is most likely to be successful if individuals are dispersing into productive sites regardless of environment of origin, whereas establishment in unproductive sites is less likely. (2) Productive patches respond strongly to climate fluctuations, frequently resetting competitor densities to low levels, promoting establishment. When competitor densities are high in all patches, establishment success is low and dispersal does not evolve (Travis and Dytham 1998). We highlight an opportunity for research on how spatial variation in average habitat quality and in the magnitude of variability in quality affect patterns of gene flow in landscapes.

To conclude, faced with increasing environmental uncertainty, populations must either evolve strategies for individuals to avoid or tolerate adverse conditions or face extinction. Understanding the intrinsic capacity for populations to adapt when challenged in this way is crucial to evaluating extinction risk as variability regimes shift with global change (Rummukainen 2012). Our study is a much needed step towards understanding the evolution of two avoidance strategies: dispersal, dormancy, and their covariance, providing new insight into the environmental drivers of bet hedging. More empirical studies are needed to move towards a general, scalable knowledge of how to best manage populations and landscapes to support the evolutionary processes that prevent extinction (Derry et al. 2019). More generally, bet hedging strategies are critical to theories of biodiversity maintenance in variable environments (e.g., storage effects (Abrams et al. 2013), metacommunity archetypes (Leibold et al. 2004)) and we document the evolution of these ecologically-important strategies.

## Acknowledgments

We thank the Angert Lab, Amy Angert, Megan Szojka, and Jeannette Whitton for feedback on our experimental design, Mackenzie Urquhart-Cronish for greenhouse assistance, Kately Nikiforuk for field assistance, and the Beaty Biodiversity Museum for lending deer pelts from their collection. This project was supported by an NSERC Discovery Grant (2019-04872) to R.M.G. We declare no conflicts of interest.

