## Appendix for "Environmental variability and the evolution of bet-hedging strategies"

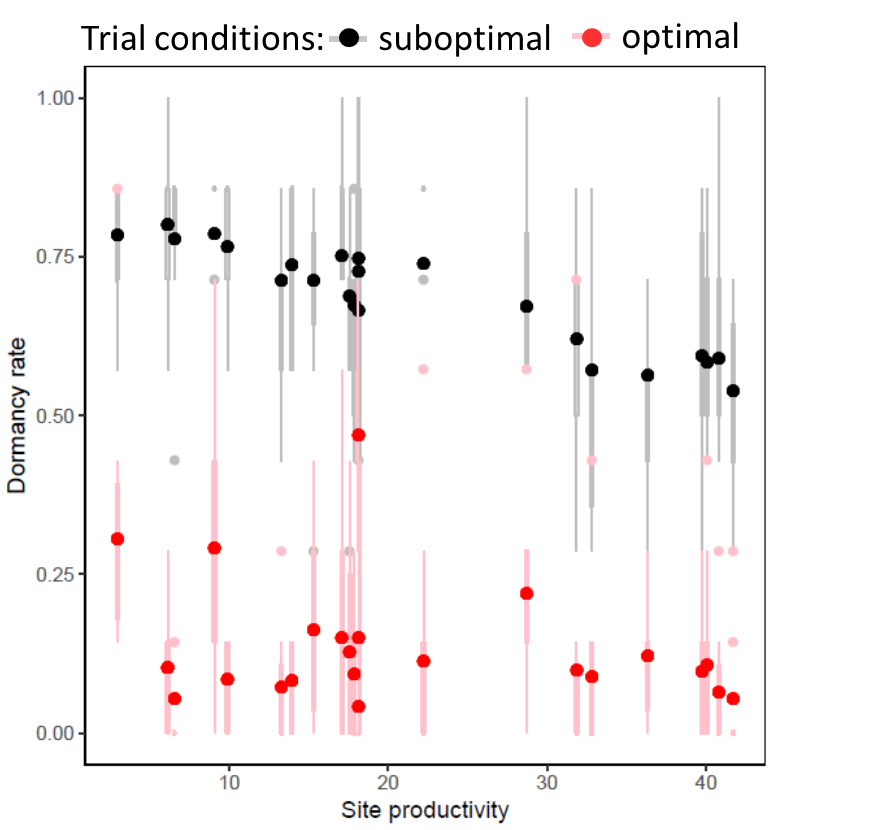

**Figure S1.** Dormancy rates were more sensitive to germination trial conditions when populations originated from low productivity sites. Points are the fitted population averages from a glme model, with boxplot silhouettes showing the distribution of data within populations. Black points are fitted population averages of dormancy rates when the germination trial was performed in suboptimal conditions (shown in Fig. 2A (i)) whereas red points are in optimal conditions.
